# Subpopulations of corticotropin-releasing factor containing neurons and internal circuits in the chicken central extended amygdala

**DOI:** 10.1101/2023.06.08.544245

**Authors:** Alessandra Pross, Alek H. Metwalli, Antonio Abellán, Ester Desfilis, Loreta Medina

**Author notes:** Serra Húnter fellow (Ester Desfilis, Loreta Medina). AP processed most of the material and analyzed it, as part of her Ph.D. research project. These authors contributed equally as supervisors of the project. Correspondence to: Loreta Medina, Ph.D., Departament de Medicina Experimental, Facultat de Medicina, Universitat de Lleida, Edifici Biomedicina-Bloc 2, IRBLleida, Avda. Alcalde Rovira Roure 80, Lleida 25198, Catalunya, Spain. **Co-author’s emails (in alphabetical order):** Antonio Abellán, Alek H. Metwalli, Alessandra Pross, Ester Desfilis. **Ethical Statement:** We declare that we adhere to the ethical and integrity policies of the journal regarding research. **Conflict of Interest:** The authors declare that the research was conducted in the absence of any commercial or financial relationships that could be construed as a potential conflict of interest. **Availability of data:** The most relevant data that support the findings of this study are included in the figures of this paper. **Funding:** Funded by grants from the European Union’s Horizon 2020 research and innovation programme under the Marie Skłodowska-Curie grant agreement No. 812777 (H2020-MSCA-ITN-2018-812777) and the Spanish Ministerio de Ciencia e Innovación (Agencia Estatal de Investigación, Grant No. PID2019-108725RB-100). AHM and AP have contracts as Early-Stage Researchers paid by the H2020-MSCA-ITN-2018-812777 project.

## Abstract

In mammals, the central extended amygdala is critical for regulation of the stress response. This regulation is extremely complex, involving multiple subpopulations of GABAergic neurons and complex networks of internal and external connections. Two neuron subpopulations expressing corticotropin-releasing factor (CRF), located in the central amygdala and the lateral bed nucleus of the stria terminalis (BSTL), play key roles in the long-term components of fear learning and in sustained fear responses akin to anxiety. Very little is known on the regulation of stress by the amygdala in non-mammals, hindering efforts for trying to improve animal welfare. In birds, one of the major problems relates to the high evolutionary divergence of the telencephalon, where the amygdala is located. In the present study, we aimed to investigate the presence of CRF neurons of the central extended amygdala in chicken and the local connections within this region. We found two major subpopulations of CRF cells in BSTL and the medial capsular central amygdala of chicken. Based on multiple labeling of CRF mRNA with different developmental transcription factors, all CRF neurons seem to originate within the telencephalon since they express Foxg1, and there are two subtypes with different embryonic origins that express Islet1 or Pax6. In addition, we demonstrated direct projections from Pax6 cells of the capsular central amygdala to BSTL and the oval central amygdala. We also found projections from Islet1 cells of the oval central amygdala to BSTL, which may constitute an indirect pathway for regulation of BSTL output cells. Part of these projections may be mediated by CRF cells, in agreement with expression of CRF receptors in both Ceov and BSTL. Our results show a complex organization of the central extended amygdala in chicken, and open new venues for studying how different cells and circuits regulate stress in these animals.

## Introduction

Regulation of the stress response by the central extended amygdala is extremely complex in mammals, involving several subdivisions, multiple subpopulations of GABAergic neurons and complex networks of internal and external connections (Ulrich-Lai and Herman, 2009; Calhoon and Tye, 2015; Zhang et al., 2021). Neurons expressing corticotropin-releasing factor (CRF) play a key role in regulation of sustained fear responses and memory (Davis et al., 2010; Gafford et al., 2012; Gafford and Ressler, 2015). In mammals, two major subpopulations of CRF neurons are found in the central extended amygdala, one located in the dorsal or oval subdivision of the lateral bed nucleus of the stria terminalis (BSTL), and another one located in the lateral subdivision of the central amygdala (Merchenthaler et al., 1982; Swanson et al., 1983; Moga and Gray, 1985; Moga et al., 1989; Shimada et al., 1989; Pitts et al., 2009; Davis et al., 2010; Gafford and Ressler, 2015; McCullough et al., 2018; Kovner et al., 2019; Ortiz-Juza et al., 2021).

However, smaller subsets of CRF cells are also found in other parts of the central extended amygdala, including the medial central amygdala and the ventral BSTL (Shimada et al., 1989; Kono et al., 2017; McCullough et al., 2018; Ortiz-Juza et al., 2021). In the central amygdala and BSTL, some of the CRF cells are segregated from other peptidergic subpopulations, but there is a subset of CRF cells that co-contain somatostatin (SST), tachykinin 2, and neurotensin (Shimada et al., 1989; McCullough et al., 2018; Kovner et al., 2019), raising questions of the specific connections and functions of these different subtypes.

CRF cells of the lateral central amygdala have inhibitory projections to the BSTL (Partridge et al., 2016; Asok et al., 2018) and to autonomic centers in the brainstem (Moga and Gray, 1985). When the projections to BSTL are silenced by way of optogenetics, the long-lasting components of fear memory are disrupted (Asok et al., 2018). In particular, these projections appear to be involved in contextually conditioned fear memories (Asok et al., 2018). The projections from CRF cells of the central amygdala target both CRF and non-CRF cells of BSTL, but chronic stress appears to enhance contacts with CRF cells (Partridge et al., 2016). The latter are involved in promoting sustained contextualized fear responses, akin to anxiety (Davis et al., 2010; Gafford et al., 2012; Gafford and Ressler, 2015).

Regulation of the fear/stress response by the extended amygdala in non-mammals is still poorly understood. In birds, this is in part due to the high divergence of the telencephalon in evolution, which has made it challenging to identify the amygdala in sauropsids (reviewed by Reiner et al., 2004; Martínez-García et al., 2007; Kuenzel et al., 2011; Medina et al., 2011, 2017, 2023). The avian BSTL was identified in the pallidal territory of the subpallium, adjacent to the lateral ventricle (Aste et al., 1998; Puelles et al., 2000; Reiner et al., 2004; Kuenzel et al., 2011). This nucleus has descending projections to hypothalamic and brainstem centers similar to those of mammalian BSTL, involved in regulation of the endocrine, autonomic and behavioral aspects of the stress response (Atoji et al., 2006; Hanics et al., 2017). Like that of mammals, the avian BSTL was also found to be involved in stress (Nagarajan et al., 2014).

However, identification of the central amygdala in birds was not possible until recently. Progress was achieved thanks to the use of an evolutionary-developmental biology approach, which is highly powerful for identification of homologies (discussed by Medina et al., 2023). Using this approach, we compared the cells of the central extended amygdala of mouse, chicken and zebra finch using a combination of developmental regulatory transcription factors (including Pax6, Islet1 and Nkx2.1) and mRNA of different neuropeptides (such as pro-enkephalin [pENK] and SST; Vicario et al., 2014, 2017; Pross et al., 2022). Similar cell populations were found in mouse, chicken and zebra finch (Bupesh et al., 2011; Vicario et al., 2014, 2017), and also appear to be present in the central amygdala of turtles (Moreno et al., 2010), suggesting that these cells are homologous (discussed by Medina et al., 2017, 2023). Based on migration assays in mouse and chicken, Pax6 cells of the central amygdala were shown to originate in the dorsal striatal embryonic domain, Islet1 cells derive from the ventral striatal embryonic domain and Nkx2.1 originate mostly in the pallidal domain, with a minor contribution of the preoptic domain (mouse: Waclaw et al., 2010; Bupesh et al., 2011, 2014; chicken: Cobos et al., 2001; Vicario et al., 2015). By multiple fluorescence labeling, many pENK cells of the central amygdala in chicken (about half in some subdivisions) coexpress Pax6 and, thus, have a dorsal striatal origin (Pross et al., 2022), being similar to ENK cells of the mouse central amygdala (Bupesh et al., 2011; Medina et al., 2023). In contrast, the majority of SST cells of the central amygdala of chicken coexpress Nkx2.1 and appear to primarily originate in the pallidal embryonic division (Pross et al., 2022), and seem to be homologous to SST cells of the mouse central amygdala (Bupesh et al., 2011; Medina et al., 2023). In addition to ENK and SST cells, like in mammals, the chicken central amygdala and the BSTL also contain subpopulations of CRF cells (Richard et al., 2004; Metwalli et al., 2023). During development, those of BSTL appear earlier than the ones in the central amygdala (Metwalli et al., 2023). The presence of these CRF cell populations raises questions on their specific embryonic origin, their connections, and their function(s). The aim of the present study was to investigate the putative embryonic origin of CRF cells of the chicken central extended amygdala by analyzing coexpression of CRF mRNA with different developmental regulatory transcription factors. Moreover, we studied local connections within the central extended amygdala of chicken to investigate if the chicken central amygdala subdivisions project directly and/or indirectly to BSTL, paying special attention to the subdivisions containing CRF cells and/or CRF receptors.

## Material and Methods

### Animals

Fertilized eggs of domestic chicks (*Gallus gallus domesticus*; White Leghorn strain) were obtained from a commercial poultry hatchery (Granja Santa Isabel, Cordoba, Spain; Authorization ES140210000002), and were incubated at 37.5 °C and 55-60% relative humidity, with automatic egg turning. The first day of incubation was considered embryonic day 0 (E0). For analysis of CRF and CRF receptor 2 expressions, we used 12 embryos of 18 incubation days (E18, equivalent to Hamburger and Hamilton’s stages HH44). For the tract tracing studies, we used 48 embryos (E18-E20), plus 4 one-day old (P1) posthatchlings, of both sexes. All animals were treated according to the regulations and laws of the European Union (Directive 2010/63/EU) and the Spanish Government (Royal Decrees 53/2013 and 118/2021) for care and handling of animals in research. The protocols used were approved by the Committees of Ethics for Animal Experimentation and Biosecurity of the University of Lleida (reference no. CEEA 08-02/19), as well as that of the Catalonian Government (reference no. CEA/9960_MR1/P3/1 for embryos, and CEA/9960_MR1/P4/1 for post-hatchlings).

### Tissue collection and fixation

For tissue collection and fixation, we followed the protocol previously described by Pross et al. (2022). Animals were anaesthetized first with Halothane (2-Bromo-2-Chloro-1,1,1-trifluoroethane, Sigma-Aldrich, Germany; 1/1000 v/v), followed by a euthanasic dose of Dolethal (100 mg/kg of sodium pentobarbital; intraperitoneal). After this, animals used for gene expression were transcardially perfused with saline followed by phosphate-buffered (PB) 4% paraformaldehyde (PFA 4%, pH 7.4, PB 0.1M). After dissection and post-fixation (24 hours at 4°C), brains were sectioned (100 μm-thick) in a coronal plane using a vibratome (Leica VT 1000S). All brain sections were maintained at 4°C (for short storage) or at −20°C (for longer storage) in hybridization buffer, until being processed as described below. Animals used for tract-tracing studies were decapitated after anesthesia, and the brains dissected and sectioned in the frontal plane using a matrix, obtaining 2-4 mm thick slices, which were placed in artificial cerebrospinal fluid supplemented with carbogen, as explained below.

### Gene expression procedures

The procedures were based on those previously settled and described by Metwalli et al. (2022, 2023) and Pross et al. (2022) for chromogenic and indirect fluorescent in situ hybridization.

### Riboprobe preparation

The riboprobes were synthetized from 2 clones that were purchased from Genscript or obtained by BBSRC ChickEST Database (Boardman et al., 2002), distributed by Source BioScience (see Table 1). For CRFR2 riboprobe synthesis, we used a standard procedure previously described (Pross et al., 2022; Metwalli et al., 2023), but for synthesizing CRF riboprobe we followed a slightly different procedure. The vector containing the CRF sequence possessed the T7 promotor to synthesize the sense RNA (forward) but not the one to produce the antisense (reverse), which is needed to study mRNA expression. To solve this problem, we designed a longer reverse primer that included the T3 promotor sequence plus a small piece of the CRF gene. The sequence of the reverse primer was: 5’GATATTAACCCTCACTAAAGGGAACCTTCCGATGATTTCCATCAGTTTC 3’ The size differences between both primers were solved by using two annealing temperatures (51 °C for 10 cycles and 56°C for 20 cycles) during the amplification and synthesis (as explained by Metwalli et al., 2023).

**Table 1:** Genes and cDNAs employed for in situ hybridization.

| **Gene name** | **Gene accession number** | **EST (expressed sequenced tag) code** | **Size (bp)** | **Obtained from** |
| --- | --- | --- | --- | --- |
| Chicken Corticotropin releasing factor (*Crf*) | NM_001123031.1 |  | 165-668 | GenScript (clone ID OGa24568) |
| Chicken Corticotropin releasing factor receptor 2 (*Crfr2*) * | NM_204454.1 | ChEST880J1 | 1–932 | Source BioScience |

\* The *Crfr2* was previously used by Vicario et al. (2014, 2021 corrigendum).

The digoxigenin-labeled antisense riboprobe was synthetized using the DIG RNA Labeling mix with the T3 RNA polymerase (Roche Diagnostics, cat. no. 11 031 171 001) following the manufacturer’s instructions. Following this, the riboprobe was purified following standard sequential steps that consisted in adding a cold solution containing 100μl TE buffer (10mM Tris-HCl pH8, 1mM ethylenediaminetetraacetic acid [EDTA]), 5μl of LiCl 8mM and 300μl Ethanol (EtOH) 100%, followed by keeping the solution at −20°C overnight to precipitate the riboprobe RNA. After, the solution was centrifuged for 30 minutes at 14.000g at 4°C, then the supernatant was discarded, and 200μl of EtOH 70% at 4°C were added to wash the pellet, which was followed by another centrifugation for 10 minutes. The resulting pellet was dried, then resuspended adding a solution 1:1 of H2O and Formamide (Fisher), and stored it at −20°C.

### Fluorescent in situ hybridization combined with single and double immunofluorescence

To study mRNA expression of the CRF and CRFR2 genes, parallel series of brain sections were processed for indirect fluorescent in situ hybridization, followed by single or double immunofluorescence, as described previously (Metwalli et al., 2022; Pross et al., 2022).

Free floating sections were prehybridized in the prehybridization buffer for 2 hours at 58°C and then hybridized in the hybridization buffer, containing the riboprobe overnight at 63°C (0,5-1µg/ml depending on the probe). The composition of buffers is described in Metwalli et al. (2022, 2023). For the fluorescent in situ hybridization, CRF was used at 0,4μg/ml, while CRFR2 was used at concentration of 0,8μg/ml.

After hybridization, the sections were thoroughly washed with the prehybridization buffer for 30 min at 58°C, followed with a wash in sodium-citrate buffer (SSC; 0.2M; pH 7.5) for 10 minutes at room temperature. Next, sections were treated with the peroxidase deactivation buffer, followed by washes in TNT (10% Tris, pH 8.0, 0.1 M; 0.9% NaCl; 0.05% Tween-20) for 15 minutes. Subsequently, the sections were treated with the blocking buffer (2% blocking reagent and 20% sheep serum in TNT) for 2h under gentle shaking. After blocking, sections were incubated in anti-DIG POD (sheep anti-digoxigenin peroxidase conjugated antibody, RRID AB_514500; Roche, diluted 1:200 in the blocking buffer) overnight, under gentle shaking, at 4°C in the dark. The next day, samples were thoroughly rinsed with the TNT buffer, then treated with the TSA/Cy3 working solution (tyramide conjugated to Cy3, Perkin Elmer, USA; diluted 1:50 and freshly prepared before the reaction) for 10 minutes in the dark. Reaction was stopped with a thorough wash with 0.1M Tris buffer. Sections were then processed for immunofluorescence.

We performed triple labelling to study the coexpression of the CRF peptide mRNA with the transcription factors Pax6, Islet1, Nkx2.1 and/or Foxg1 (detected by immunofluorescence), while the double labelling procedure was done to assess if CRFR2 expressing neurons in the Ceov co-express Islet1. For the triple labelling experiment, one series of parallel brain sections was used to detect CRF, followed by immunofluorescence for Pax6 combined with either Nkx2.1 or Foxg1, while the other series was processed for CRF combined with immunofluorescence for Islet1 and Nkx2.1.

The primary and secondary antibodies used for immunofluorescence are listed in Table 2.

**Table 2:**
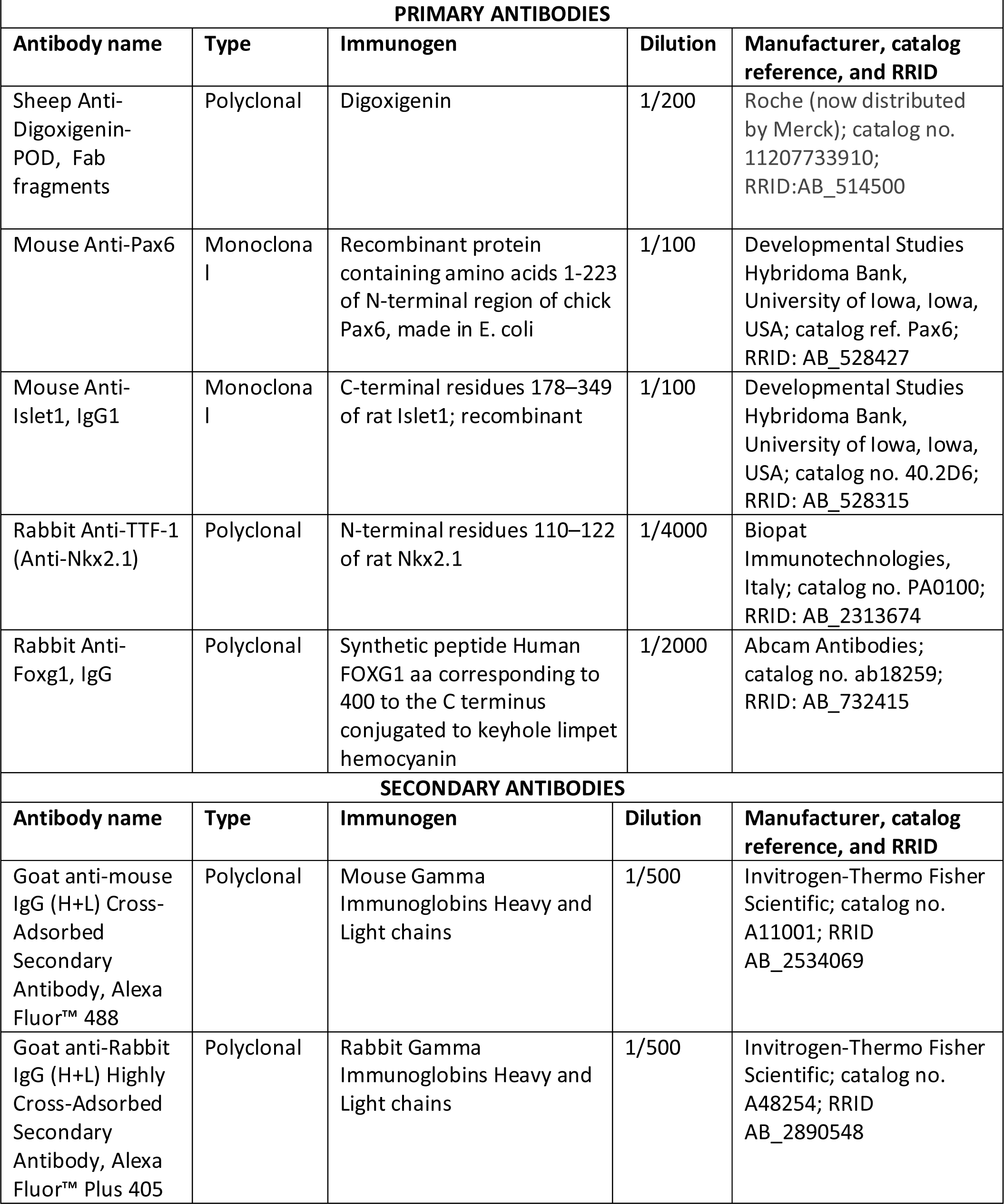
Primary antibodies.

All antibodies were diluted in phosphate-buffered saline (PBS, 0.1M, pH 7.4), containing Triton X-100 (PBST). The specificity of primary antibodies has been checked by the respective manufacturers (details in Table 2). Regarding antibodies produced raised against non-chicken proteins (Islet1, Nkx2.1, Foxg1) but used here to detect these proteins in chicken brain tissue, they were used in previous studies and demonstrate that:

- In the forebrain, the anti-Islet 1 produces a staining pattern identical (Pross et al., 2022) to that observed for the mRNA expression, using a riboprobe specific for chicken Islet1, and highly similar to that obtained with other specific antibodies (Vicario et al., 2014, 2015).
- Similarly, staining in the chicken brain with the anti-Nkx2.1 antiserum is identical to that of the mRNA signal of chicken Nkx2.1 (Abellán and Medina, 2009). The specificity of the anti-Nkx2.1 has also been demonstrated in other sauropsids (turtles) by Western blot (Moreno et al., 2010).
- Regarding Foxg1, in the brain this antibody produces a staining pattern (Metwalli et al., 2022) identical to that observed for the mRNA expression using a riboprobe for chicken Foxg1 (Crossley et al., 2001).

At the end, the sections were mounted and cover slipped using an antifading mounting medium (Vectashield Hardset Antifade mounting medium, Vector Laboratories Ltd., UK).

### Tract tracing experiments

We carried out tract-tracing experiments in organotypic cultures of chicken brain slices, using biocytin as tracer, following the protocol described by Ahumada-Galleguillos et al. (2015). Chickens are precocial animals, and the late embryonic/neohatchling brain is already quite mature, but smaller than that of adult animals. As a consequence, thick frontal slices of the brain of these young animals at the level of the anterior commissure include all areas of interest (i.e. all subdivisions of the extended amygdala) and also preserve fiber tracts running across them, allowing the study of connections between these areas in this ex-vivo condition.

First, the animals were anaesthetized with Halothane, as previously described. Then, they were decapitated and the brain was rapidly removed from the skull. The brain was immediately submerged in a cooled (4°C) Krebs solution (240 mM sucrose, 3 mM KCl, 3 mM MgCl2, 23 mM NaHCO3, 1.2 mM NaH2PO4, 11 mM D-glucose), continuously bubbled with carbogen (95% O2, 5% CO2) for oxygenation during 10 minutes.

The brains were then placed in a matrix and sliced in the frontal plane at 2-4 mm thickness. Sections containing the level of the anterior commissure were collected and used to study the internal connectivity of the central extended amygdala. The slices were placed in a well containing artificial cerebrospinal fluid (ACSF; 119 mM NaCl, 2.5 mM KCl, 1.3 mM Mg2SO4, 1.0 mM NaH2PO4, 26.2 mM NaHCO3, 11 mM D-glucose, 2.5 mM CaCl2), continuously supplemented with carbogen, and then they were placed on an insert (Millicell cell culture insert) inside a well plate containing ASCF. Subsequently, a small biocytin crystal (Sigma-Aldrich, St. Louis, MO; B4261) was deposited on the surface of the slice, selecting different targets (mostly BSTL and Ceov). The crystals were manually deposited under visual control using a stereotaxic microscope, with a fine glass micropipette (sealed in the back to avoid liquid filling by capillarity when placing the crystal). Then, brain slices were maintained in culture with ACSF continuously supplemented with carbogen, at 20°C, for a period of 4-6 hours, to allow the biocytin tracer to be caught by cell bodies and terminals in the area of the crystal deposit and to spread through the neuronal axons. After that time, the slices were removed from the insert, washed 3 times with PBS and then fixed in PFA 4% for 2/4 hours, depending on the thickness of the tissue (1 hour for each mm). Finally, the sections were cryoprotected in 30% sucrose, and then they were frozen for subsequent use.

The slices were re-sectioned at 100μm, using a freezing microtome. Each section was placed in a different well, using a 24 multi-well plate, to facilitate following the correct slice sequence. We obtained three different series of parallel sections and used each series to do single or multiple staining of different markers. In most cases, the first series was processed using the chromogenic protocol to allow the visualization of the tracer and evaluate the precision of the deposit. For this, we first inhibited the endogenous peroxidase activity by incubating in 1% H2O2 and 2% methanol in PBS for 30 minutes. Color reaction was obtained by incubation with Vectastain Elite ABC Kit (PK-6100, Vector Laboratories), followed by diaminobenzidine staining (SIGMAFAST, 3,3’ – Diaminobenzidine tablets, Sigma-Aldrich; diluted following the manufacturer instructions). Finally, the sections were rinsed and mounted on gelatinized glasses; then they were dehydrated and cover slipped with Permount (Fisher Scientific).

The other two series of sections of selected samples (those in which the biocytin crystal was correctly placed and had a small size) were processed for double or triple fluorescence: one series was stained to visualize Pax6, Nkx2.1 and biocytin, while the other one was processed for Islet1, Nkx2.1 and biocytin, following the protocol described previously for double immunofluorescence (Pross et al., 2022). Briefly, after blocking the unspecific binding using the blocking reagent, the sections were incubated in primary antibodies (explained above and in Table 2; Nkx2.1 and Pax6; or Nkx2.1 and Islet1) for about 60-64 hours at 4°C. Following this, sections were washed and incubated overnight, at 4°C, in fluorescence secondary antibodies (anti-mouse and anti-rabbit, coupled to Alexa Fluor 488 or 405, respectively) together with Cy3-streptavidin conjugated (1/500; iFluor Cy3-streptavidin AATBioquest), diluted in PBST. Finally, after a few washes in PBS the sections were mounted, and cover slipped using an antifading mounting medium (Vectashield Hardset Antifade mounting medium).

### Digital photographs and figures

Digital microphotographs from chromogenic experiments were taken on a Leica microscope (DMR HC, Leica Microsystems GmbH) equipped with a Zeiss Axiovision Digital Camera (Carl Zeiss, Germany), using 1.6X and 5X magnification objectives. Serial images from fluorescent material were taken with a confocal microscope (Olympus FV1000, Olympus Corporation, Japan) using 10X, 20X and 40X objectives; Z-series stacks were taken at 2 μm-steps to allow analysis of co-expression. The fluorescent images were adjusted for brightness and contrast and then extracted using Olympus FV10-ASW 4.2 Viewer (Olympus Corporation). All the figures were mounted using CorelDraw 2022 (Corel Corporation, Canada).

### Nomenclature

For the chicken brain, we used the general nomenclature proposed by the Avian Brain Nomenclature Forum (Reiner et al., 2004) and the chick brain atlas (Puelles et al., 2019). For the developing chicken brain, we followed Puelles et al. (2000) and our own publications on the subject (Abellán and Medina, 2009; Abellán et al., 2009, 2010, 2014; Medina et al., 2019), while for the chicken central extended amygdala we followed our detailed description of this territory during development (Vicario et al., 2014, 2015; Metwalli et al., 2022; Pross et al., 2022).

## Results

### *CRF* mRNA expressing cells of the chicken central extended amygdala and coexpression with Nkx2.1, Islet1, Pax6 and Foxg1

We first mapped the distribution of cells expressing *CRF* mRNA in the chicken central extended amygdala, from anterior to posterior levels, by way of fluorescent in situ hybridization (Fig. 1A-D). This was facilitated by the use of multiple labeling of *CRF* with different transcription factors, which allowed distinction of major subdivisions, as described previously (Vicario et al., 2014). Two major subpopulations were found: one located medially, in the BSTL (Fig.1A-D), and another one located laterally, in the capsular part of the central amygdala (CeC, Fig. 1C,D). In the BSTL, the majority of the cells were found in the medial and intermediate subdivisions, while in the CeC the CRF cells showed a trend to be more concentrated in the medial aspect of this subdivision (CeCm), spreading a bit over and partially overlapping the dorsal aspect of the oval central nucleus (double asterisk in Fig. 1C). Scattered *CRF* expressing cells were also found in the lateral subdivision of BSTL and the peri-/post-intrapeduncular island field (pINP).

**Figure 1.**
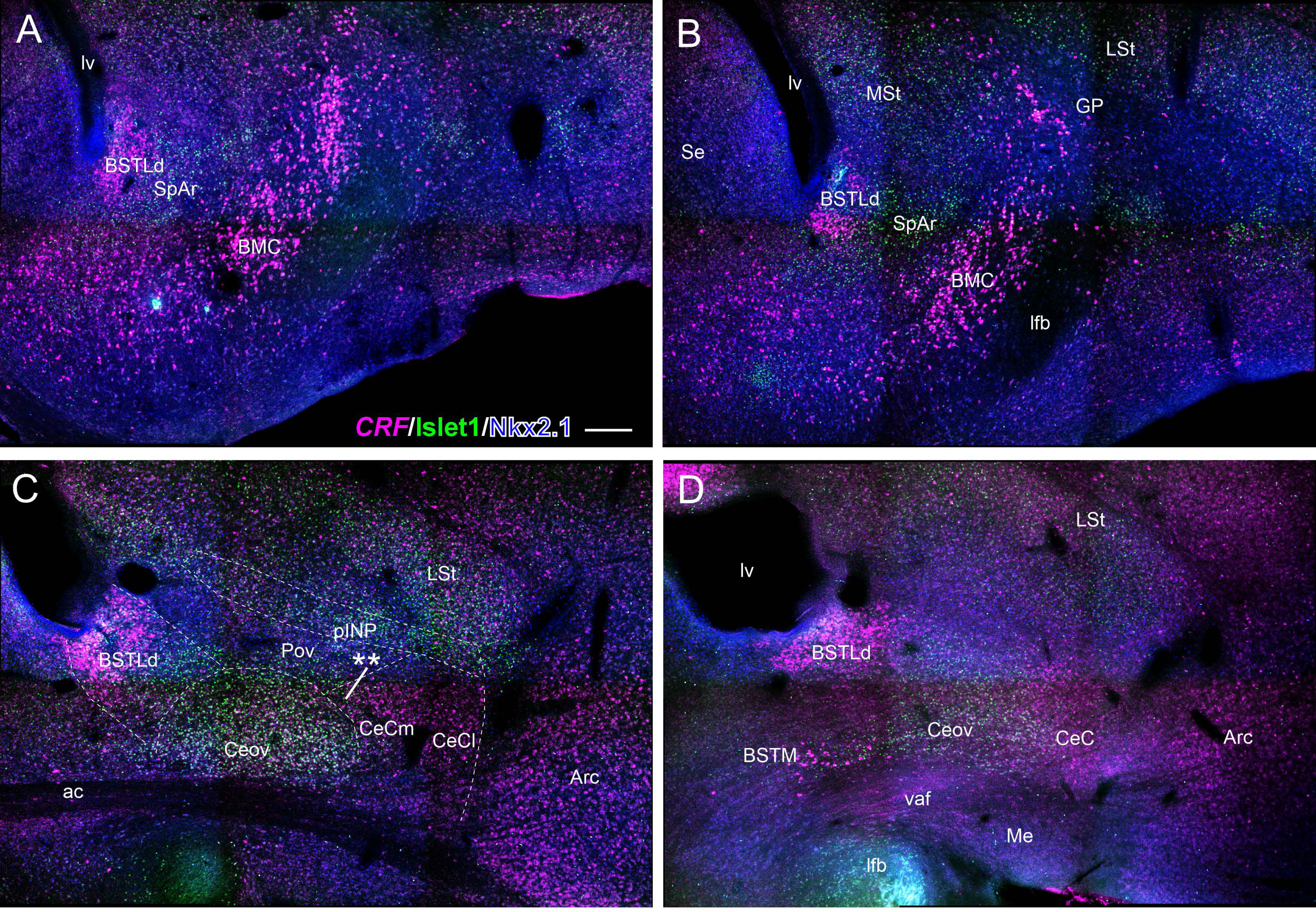
Triple fluorescence labeling of corticotropin releasing factor mRNA (*CRF*), Islet1 and Nkx2.1 in the chicken central extended amygdala at E18. A, B, C, D: Confocal microscope image mosaic (10X objective) of the subpallium, in frontal sections, at pre-commissural (A, B), commissural (C), and post-commissural levels (D). Sections were processed for indirect fluorescent in situ hybridization for *CRF* (magenta) and double immunofluorescence for Islet1 (green) and Nkx2.1 (blue). The combination with Islet1 and Nkx2.1 allowed better mapping of the exact location of the *CRF* cells (for example, the oval central amygdala [Ceov] is well delineated by its high amount of Islet1 cells). Two main groups of *CRF* expressing cells are observed in the central extended amygdala (EAce): one medial, located in the BSTL from anterior to posterior levels (mainly its dorsal part, related to EAce); and another one lateral, mainly located in the medial aspect of the capsular central amygdala (CeCm), spreading medially (** in panel C) over the oval central amygdala nucleus (Ceov). Some of the *CRF* cells also overlapped the dorsal aspect of Ceov. See text for more details. For abbreviations see list. Scale: bar in A = 250 μm (applies to A, B, C, D).

Regarding BSTL, *CRF* cells did not express the transcription factor Nkx2.1 in the main dorsolateral part of this nucleus, but a few double-labeled cells were found in the ventromedial aspect (Fig. 2A, detail in B-B’’’), which belongs to the medial extended amygdala (as part of BSTM complex) instead of the central extended amygdala (Abellán and Medina, 2009). In contrast, double labeling of *CRF* and the transcription factor Islet1 showed that the majority of the *CRF* cells in BSTL coexpressed Islet1 (Fig. 2A, detail in D-D’’). In addition, double-labeling of *CRF* and the transcription factor Pax6 showed cases of coexpression, mainly in the dorsal aspect of BSTL (Figs. 3A,B; details in 3C, and 3D-D’’’). However, many *CRF* cells did not coexpress Pax6 (Fig. 3E-E’’’). Since a subset of Pax6 cells of BSTL has been suggested to have an extratelencephalic origin (Abellán and Medina, 2009; Vicario et al., 2015), we carried out triple-labeling of *CRF* and the transcription factors Pax6 and Foxg1 (a telencephalic marker) (Fig. 4). Our results showed that all *CRF* cells of BSTL coexpressed Foxg1, meaning that they are telencephalic (Fig. 4A, detail in B-B’’’). Regarding the Pax6 cells of BSTL, we found a few cases without coexpression of Foxg1, confirming their extratelencephalic origin (Fig. 4A, pointed with empty arrows in details in B-B’’’ and C-C’’’).

**Figure 2.**
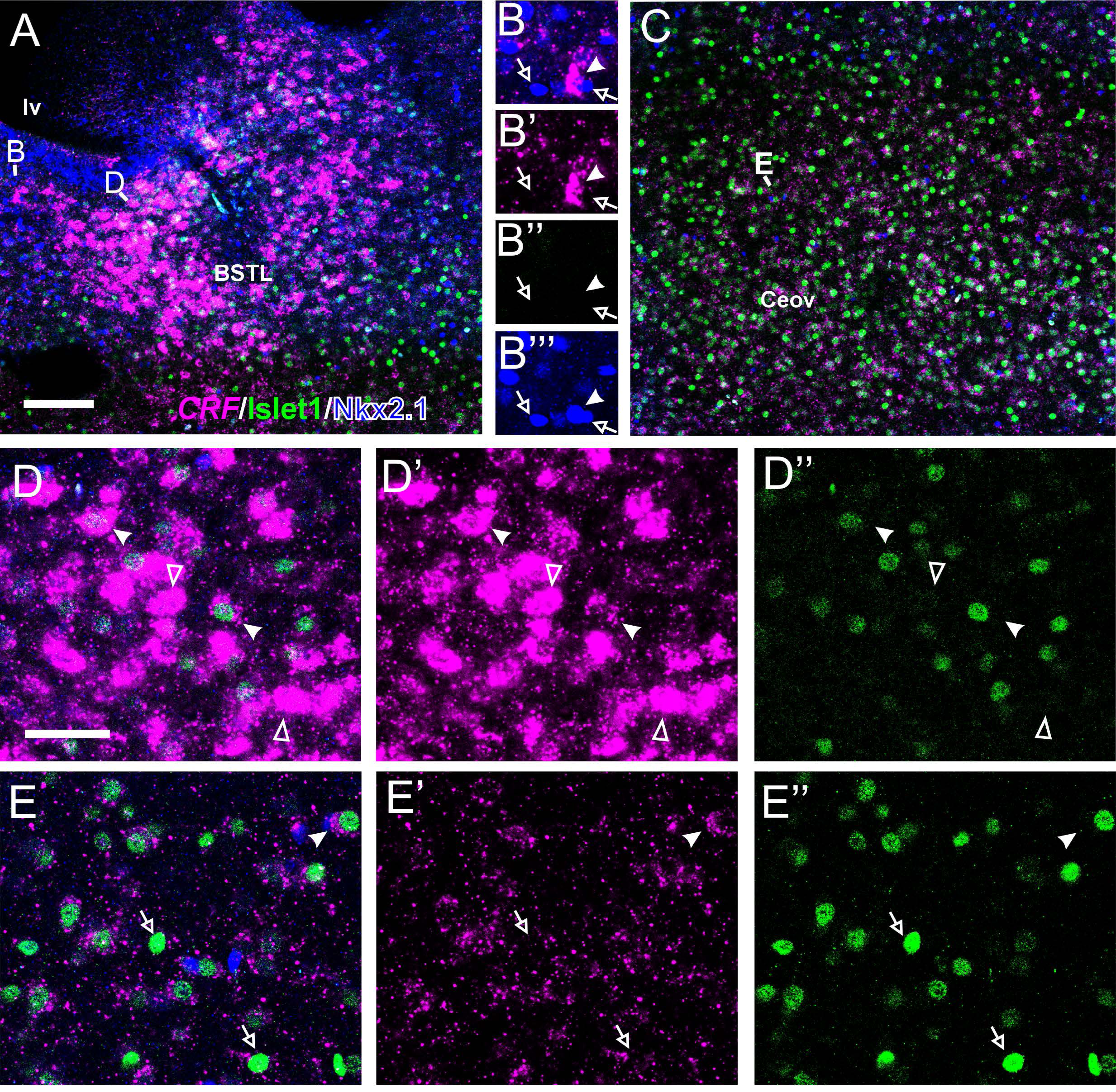
Details of triple fluorescence labeling of corticotropin releasing factor mRNA (*CRF),* Islet1 and Nkx2.1 in the chicken central extended amygdala at E18. These details are taken from the frontal section shown in Figure 1C, at the level of the anterior commissure, processed for indirect fluorescent in situ hybridization for *CRF* (magenta) and double immunofluorescence for Islet1 (green) and Nkx2.1 (blue). A, C: Details of BSTL (A) and Ceov (C) (10X objective). Note the high amount of *CRF* cells in BSTL, but the scarcity in Ceov (except a few located in its dorsal aspect). Higher magnification details of BSTL (objective 20X, only one single z level of the confocal stack) are shown in B-B’’’ (for ventral BSTL, BSTLv) and D-D’’ (for dorsal BSTL) (areas amplified indicated with the letters B and D in panel A). E-E’’ show details of the Ceov (area amplified indicated with the letter E in panel C). In B-B’’’ merged (B, with CRF in magenta and Nkx2.1 in blue), plus separate magenta (B’, CRF), green (B’’, Islet1) and blue (B’’’, Nkx2.1) channels are shown, while in D-D’’ and E-E’’ merged (D, E, with CRF in magenta and Islet1 in green), magenta (D’, E’, CRF) and green (D’’, E’’, Islet1) channels are shown. In these details, cells coexpressing *CRF* and Islet1 in D and E, and Nkx2.1 in B are pointed with a filled arrowhead, cells single labeled for *CRF* are pointed with an empty arrowhead, while cells single labeled for Islet1 or Nkx2.1 are pointed with an empty arrow (only a few examples are pointed). See text for more details. For abbreviations see list. Scale bars: A = 100μm, applies to A and C; D = 40μm applies to B, D, E.

**Figure 3.**
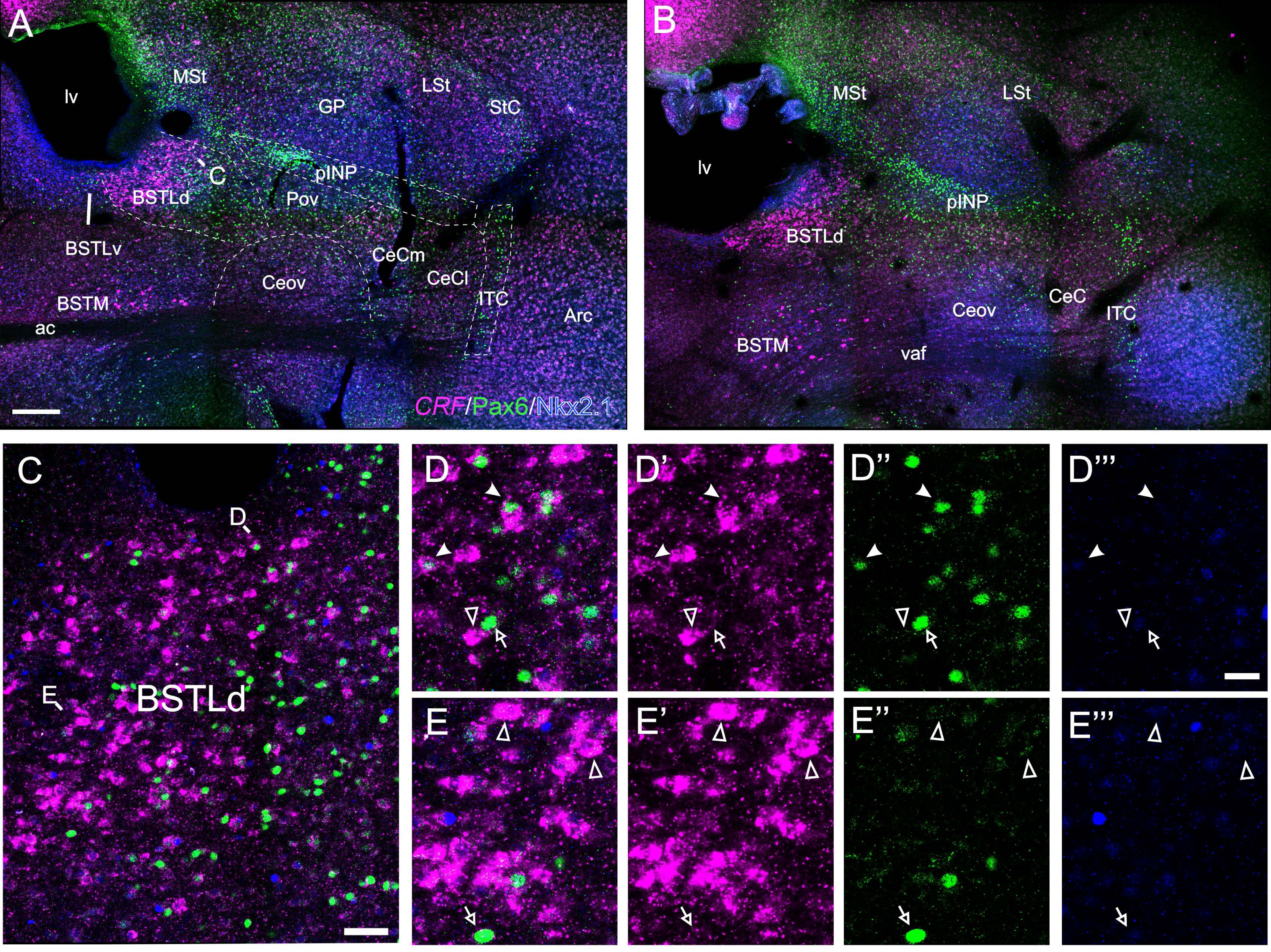
Triple fluorescence labelling of corticotropin releasing factor mRNA (*CRF*), Pax6 and Nkx2.1 in the chicken central extended amygdala at E18. A, B: Confocal microscope image mosaic (10X objective) of the subpallium in frontal sections at commissural level (A) and post-commissural level (B), processed for indirect fluorescent in situ hybridization for *CRF* (magenta,) and immunofluorescence for Pax6 (green) and Nkx2.1 (blue). Note the typical distribution of *CRF* as described in figure 1. Pax6 allows better distinction of specific subdivisions of the central extended amygdala, such as pINP and ITC. C: Detail of BSTL (10X objective) from the sections shown in panel A (area amplified is indicated with the letter C in panel A). D-D’’’, E-E’’’ show higher magnification of the areas pointed in C, with the letters D and E, respectively, in separate channels (merged in D and E, CRF in D’ and E’; Pax6 in D’’ and E’’, and Nkx2.1 in D’’’ and E’’’). In these details, cells coexpressing *CRF* and Pax6 are pointed with a filled arrowhead, cells single labeled for *CRF* are pointed with an empty arrowhead, while cells single labeled for Pax6 are pointed with an empty arrow (only a few examples are pointed). See text for more details. For abbreviations see list. Scale bars: A = 250μm applies to A, B. C = 100μm; D’’’ = 50μm (applies to D-D’’’ and E-E’’’).

**Figure 4.**
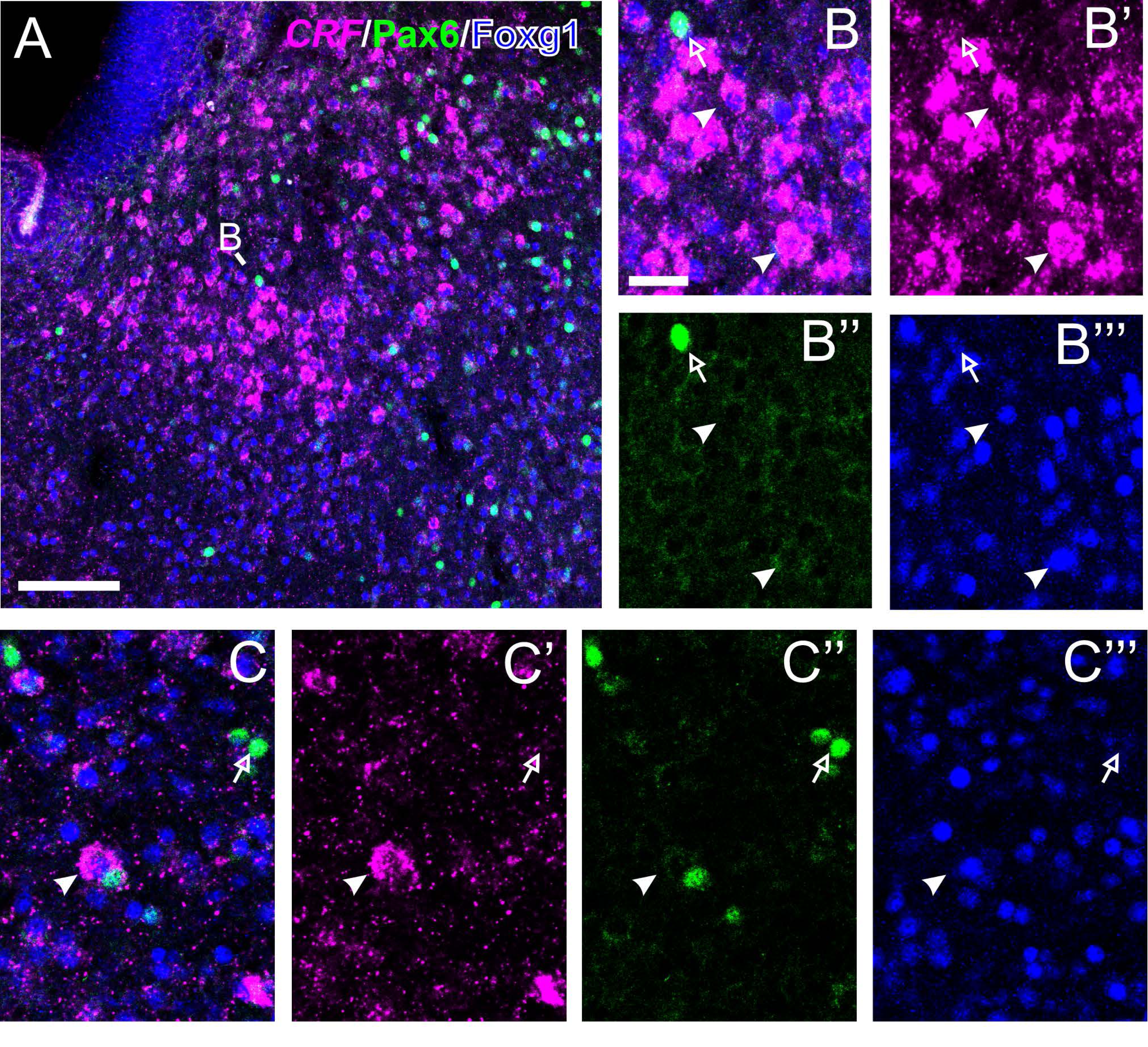
Triple fluorescence labeling of corticotropin releasing factor mRNA (*CRF*), Pax6 and Foxg1 in the chicken central extended amygdala at E18. A: Detail of the BSTL, taken from a frontal section of the chicken embryonic telencephalon at commissural level, hybridized for *CRF* (magenta), Pax6 (green) and Foxg1 (blue). B-B’’’ show a higher magnification detail of the area pointed with B in panel A (20X objective, only one single z level of the confocal stack, merged [B], plus separate magenta [B’], green [B’’] and blue [B’’’] channels are shown in separate panels). Cells coexpressing *CRF* and Foxg1 are pointed with a filled arrowhead, while cells single labeled for Pax6 are pointed with an empty arrow (only a few examples are pointed). C-C’’’ show a detail of the *CRF* cells located laterally, between Ceov and pINP, at commissural level (20X objective, only one single z level of the confocal stack, merged [C] plus separate magenta [C’], green [C’’] and blue [C’’’] channels are shown). Cells coexpressing *CRF* and Foxg1 are pointed with a filled arrowhead, while cells single labeled for Pax6 are pointed with an empty arrow (only a few examples are pointed). See text for more details. For abbreviations see list. Scale: bar in A = 100μm; B = 25 μm (applied to B-B’’’, C-C’’’).

With respect to the central amygdala, we focused on the medial aspect of CeC and its medial extension (** in Fig. 5A) over the oval central nucleus (Ceov), where the majority of the *CRF* cells locate. Our results on double-labeling of *CRF* and Pax6 showed some cases of cells coexpressing both (Fig. 5B-B’’’). In addition, we found cases of coexpresion in CeC following double-labeling of *CRF* and Islet1 (Fig. 5C-C’’’). In relation to Ceov, it is characterized by high content in Islet1 expressing cells with striatal origin (Figs. 1, 2B). This nucleus was previously thought to contain *CRF* cells, but this was amended to say that it rather contains cells expressing *CRF* receptor 2 (Vicario et al., 2014-corrigendum). This was corroborated in the present study, showing that Ceov shows expression of *CRF* receptor 2 (Fig. 6A-A’’), but is poor in *CRF*, with the apparent exception of a few cells with extremely low signal (Fig. 2E). Double-labeling of *CRF* receptor 2 and Islet1 showed that the vast majority (but not all) of the Islet1 cells of Ceov coexpress the receptor (Fig. 6A-A’). Some of the Islet1 cells of BSTL also express *CRF* receptor 2 (Fig. 6B-B’’).

**Figure 5.**
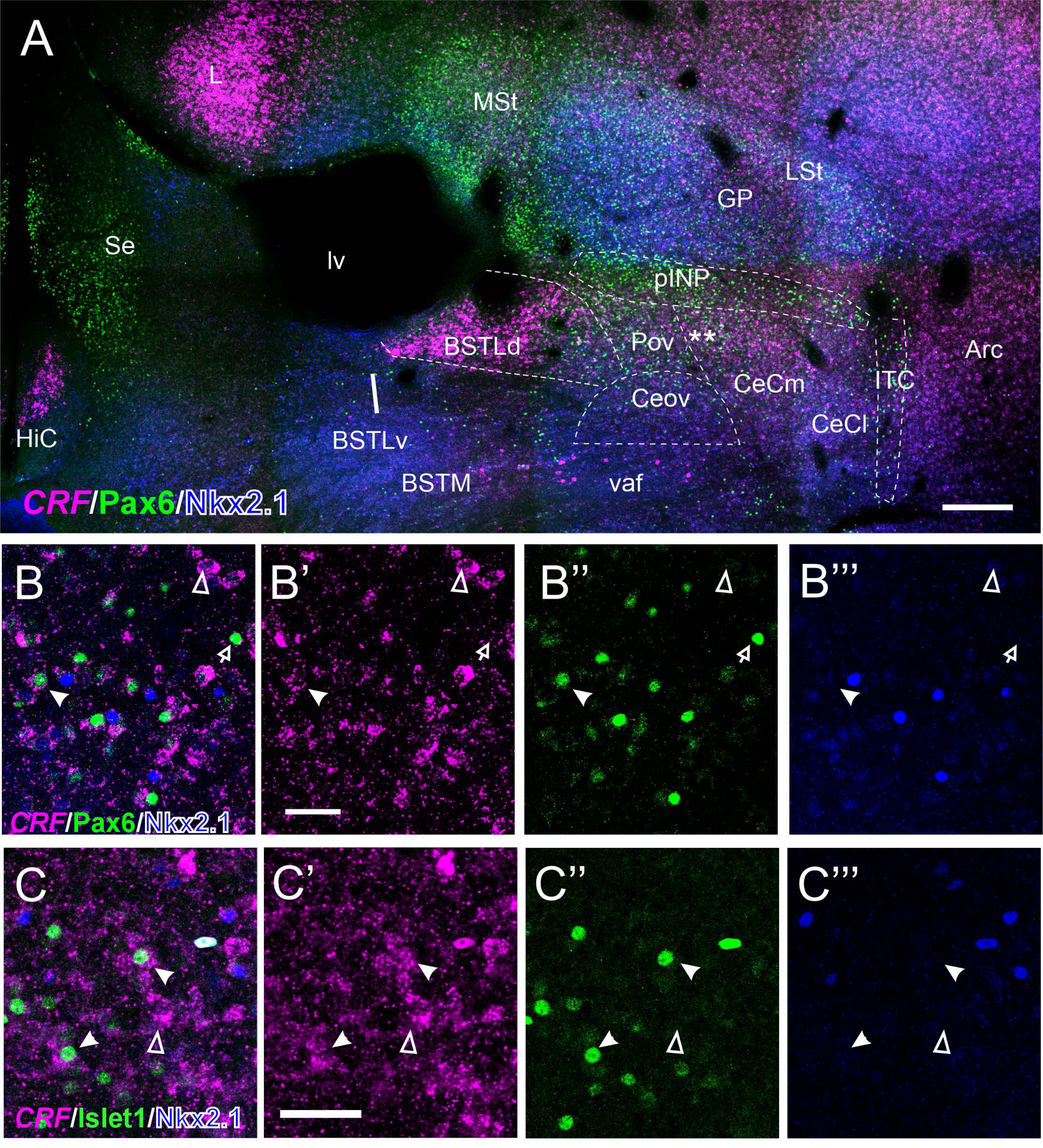
Triple fluorescence labeling of corticotropin releasing factor mRNA (*CRF*), Nkx2.1 and either Pax6 (A, B’) or Islet1 (C-C), in the chicken central extended amygdala at E18. A: Confocal microscope image mosaic (10X objective) of the subpallium in a frontal section at a post commissural level, processed for indirect fluorescent in situ hybridization for *CRF* (magenta), and double immunofluorescence for Pax6 (green) and for Nkx2.1 (blue). B-B’’’ show a high magnification detail of the area pointed with double asterisk in A, extending medially from those in CeC (20X objective, only one single z level of the confocal stack) (merged [B] plus separate magenta [B’], green [B’’] and blue [B’’’] channels are shown in separate channels). In B-B’’’, cells expressing *CRF* are pointed with empty arrowhead and cells single labeled for Pax6 are pointed with an empty arrow (only a few examples are pointed), while cells coexpressing *CRF* and Pax6 are pointed with a filled arrowhead (only very few are seen). C-C’’’ show a high magnification detail (20X objective) of the same area labeled with a double asterisk of the central amygdala, but now with triple fluorescence of *CRF* (magenta, C’), Islet1 (green, C’’) and Nkx2.1 (blue, C’’’). Cells expressing only *CRF* are pointed with empty arrowhead, cells single labeled for Islet1 are pointed with an empty arrow (only a few examples are pointed), while cells coexpressing *CRF* and Islet1 are pointed with a filled arrowhead. See text for more details. For abbreviations see list. Scale bars: A = 100μm; B’ = 50μm (applies to B-B’’’); C’ = 50μm (applies to C-C’’’).

**Figure 6.**
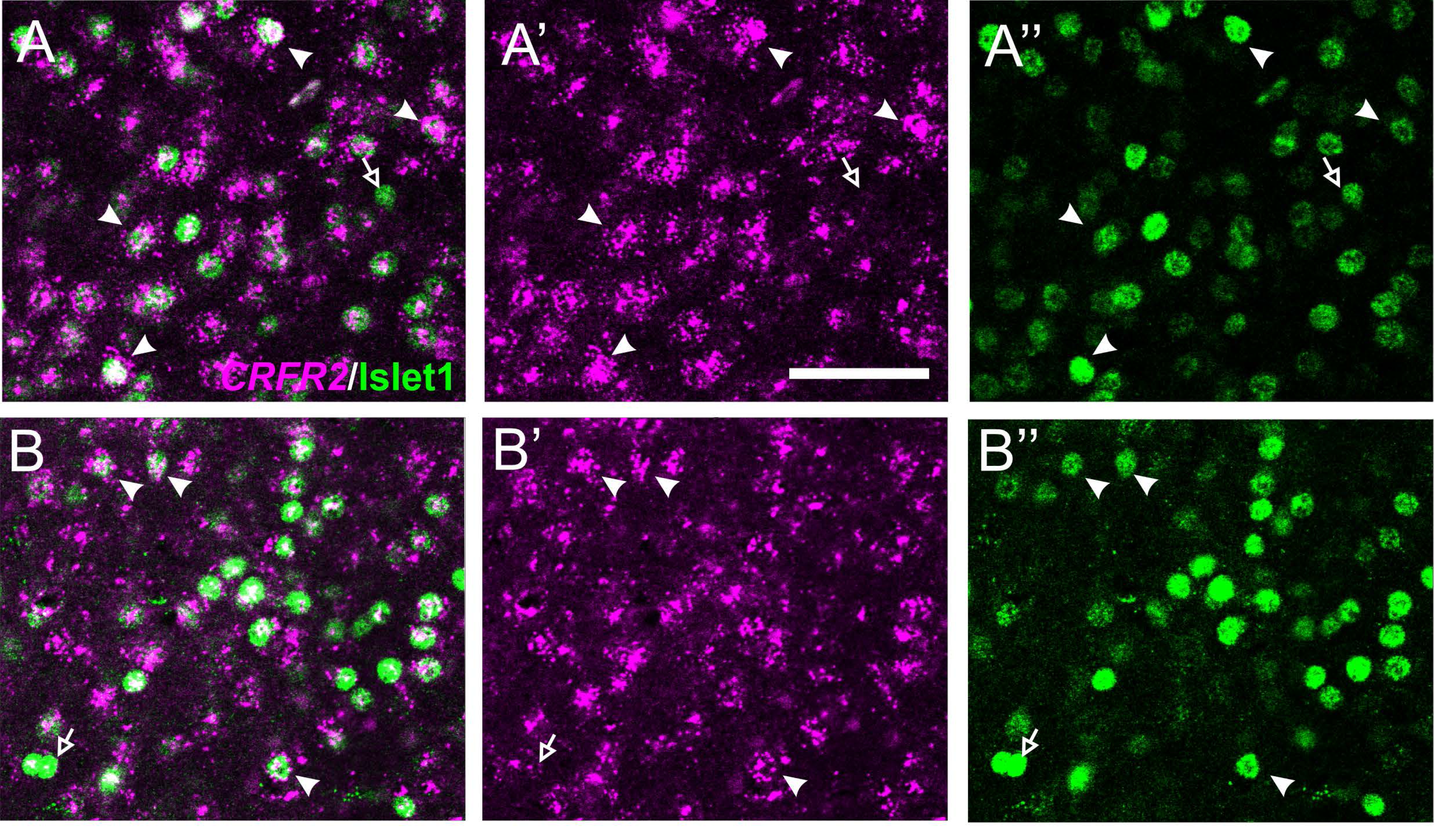
Details of double fluorescence labeling of corticotropin releasing factor receptor 2 mRNA *(CRFR2)* and Islet1 in the chicken central extended amygdala at E18. A-A’’ is a high magnification detail (objective 40X) of cells in the oval central nucleus (Ceov), while B-B’’ is a high magnification detail of cells in BSTL. *CRFR2* (magenta) was detected by fluorescent in situ hybridization, while Islet1 (green) was detected by double immunofluorescence. In these details, cells coexpressing *CRFR2* and Islet1 are pointed with a filled arrowhead, while cells single labeled for Islet1 are pointed with an empty arrow (only a few examples are pointed). See text for more details. For abbreviations see list. Scale bars: A’ = 50μm applies to A-A’’ and B-B’’.

### Connectivity between central amygdala subdivisions of chicken

To investigate whether, like in mammals, chicken BSTL receives inputs from the central amygdala, particularly from subdivisions containing *CRF* cells, we carried out tract-tracing experiments in ex-vivo conditions, applying biocytin crystals in BSTL in thick frontal brain slices containing the central extended amygdala. After application of biocytin in BSTL, retrogradely labeled cells and fibers were found in the subpallial amygdala region located between BSTL and arcopallium (Fig. 7), thus confirming previous results in pigeon and chicken (Atoji et al., 2006; Hánics et al., 2017). Here we focused our attention on the retrogradely labeled cells. Thanks to double- and triple-labeling with different transcription factors, we provide a description of the location of the cells in the different subdivisions of the central amygdala, including Ceov, pINP, and CeC (Fig. 8). A few intercalated cells (ITC, interposed between CeC and arcopallium, at the pallio-subpallial boundary) were also observed (Fig. 8C). In addition, we found cells in the medial extended amygdala (BSTM and medial amygdala; Fig. 8C). This suggests that the afferent connections of BSTL from other parts of the extended amygdala are quite similar to those found in mammals. This includes direct inputs from areas containing *CRF* cells (among other subtypes), such as CeC.

**Figure 7.**
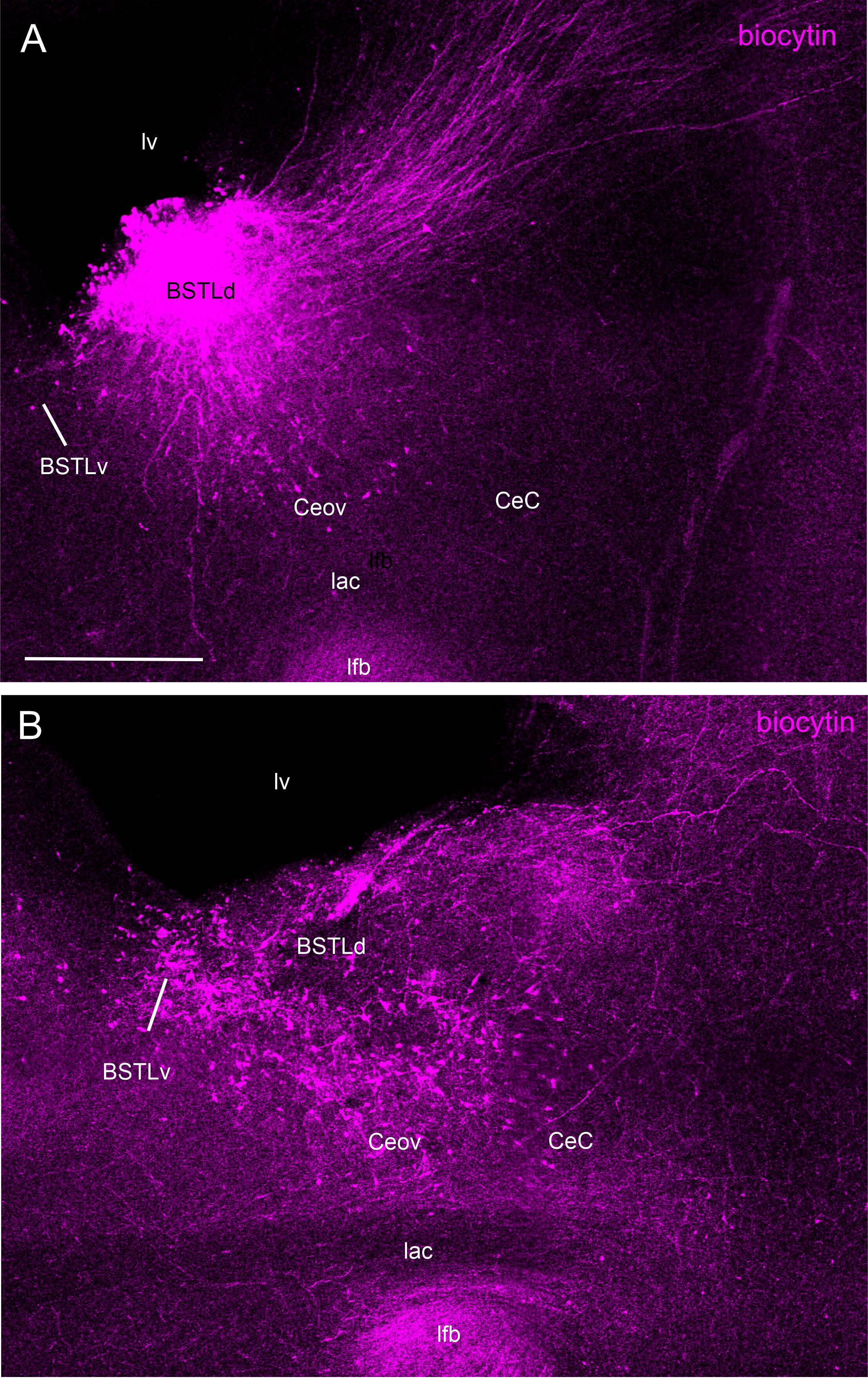
Results of a tract-tracing experiment, following deposit of biocytin crystal in a E18 chicken BSTL at the level of the anterior commissure. The biocytin was visualized in fluorescence with Cy3-streptavidin. A shows the section level of the biocytin deposit and B is a section in bit posterior to the deposit. Note the retrogradely labeled cells in the oval central nucleus (Ceov), just above the lateral branch of the anterior commissure (lac), and in the capsular central amygdala (CeC). See text for more details. For abbreviations see list. Scale bar: 500 μm.

**Figure 8.**
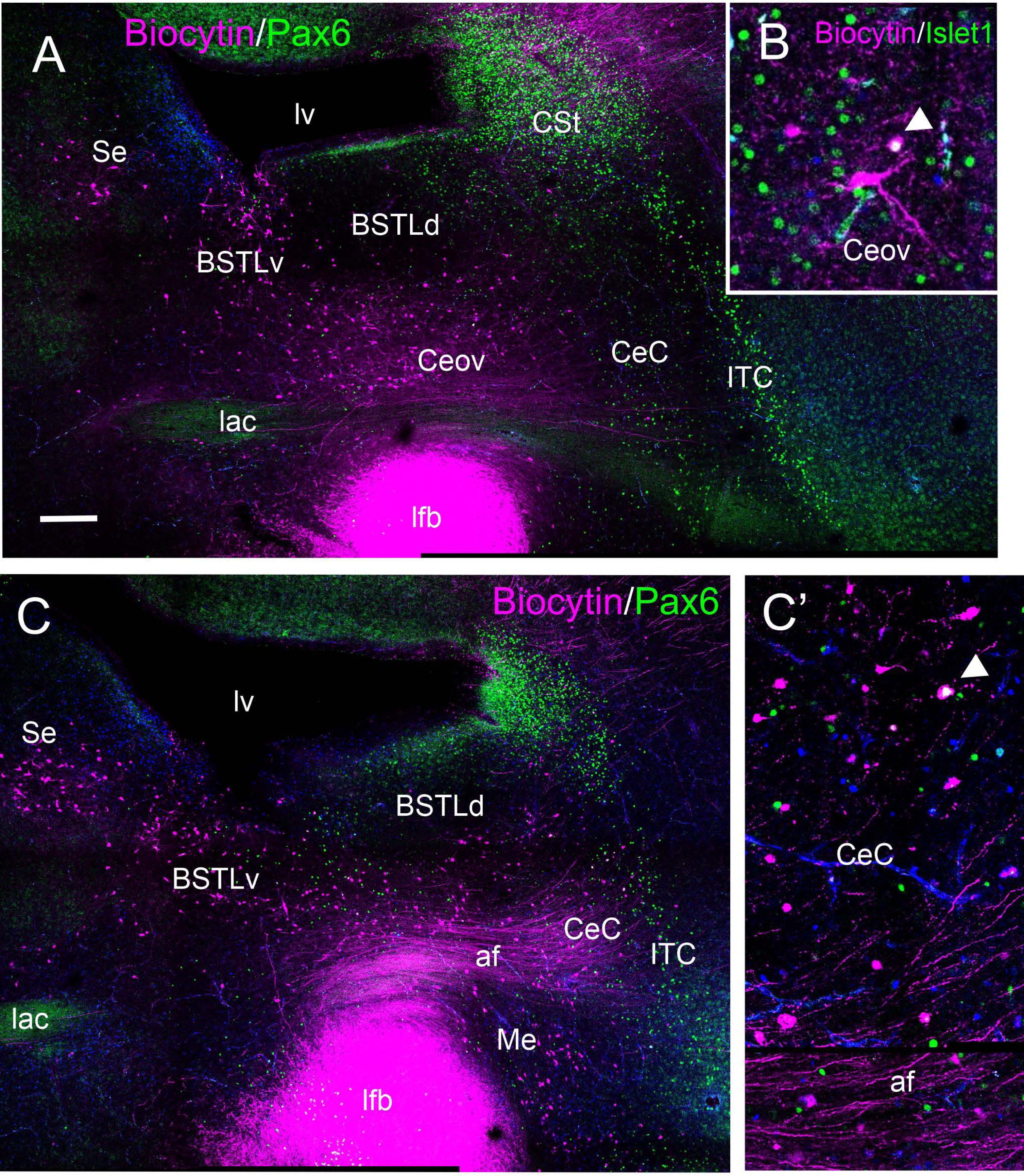
Results of a tract-tracing experiment, following deposit of biocytin crystal in a E18 chicken BSTL, at precommissural levels. Two sections of the same case are shown, at commissural (A) and postcommissural levels (C). Note the retrogradely labeled cells in different subdivisions of the central extended amygdala, including the oval central nucleus (Ceov) and the capsular central amygdala (CeC). Double labeling of biocytin and Islet1 showed coexpression in some cells of Ceov that project to BSTL (detail in B). Moreover, double labeling of biocytin with Pax6 showed coexpression in some cells of CeC that project to BSTL (detail in C’). See text for more details. For abbreviations see list. Scale bar: A = 500 μm (applies to A, C).

To get insight into the phenotype of the central amygdala neurons that project to BSTL in chicken, some cases were processed for double or triple fluorescence labeling combining biocytin (labeled with Cy3-coupled streptavidin) with immunofluorescence for Pax6 and/or Islet1, which allowed distinction of neurons with different embryonic origin and phenotype (as noted above). Our results showed a few cases of retrogradely labeled cells in CeC and pINP coexpressing Pax6 (Fig. 8C’), and a few cases of retrogradely labeled cells in Ceov coexpressing Islet1 (Fig. 8B).

Since most Islet1 cells of Ceov express *CRF* receptor 2, we also investigated the possibility that this nucleus receives inputs from CeC, which contains *CRF* cells. Thus, we carried out experiments in which biocytin was applied in Ceov. These experiments produced retrogradely labeled cells in BSTL, pINP, CeC and ITC (Fig. 9A). Some of labeled cells in CeC and ITC coexpress Pax6 (Fig. 9B,C). The results point to the existence of internal connections between subdivisions of the central amygdala, including a pathway from CeC (which contains a major subpopulation of CRF cells) to Ceov. Since Ceov projects to BSTL, the CeC-Ceov projection may be part of an indirect pathway by way of which the central amygdala can modulate BSTL, thus regulating activation of HPA axis.

**Figure 9.**
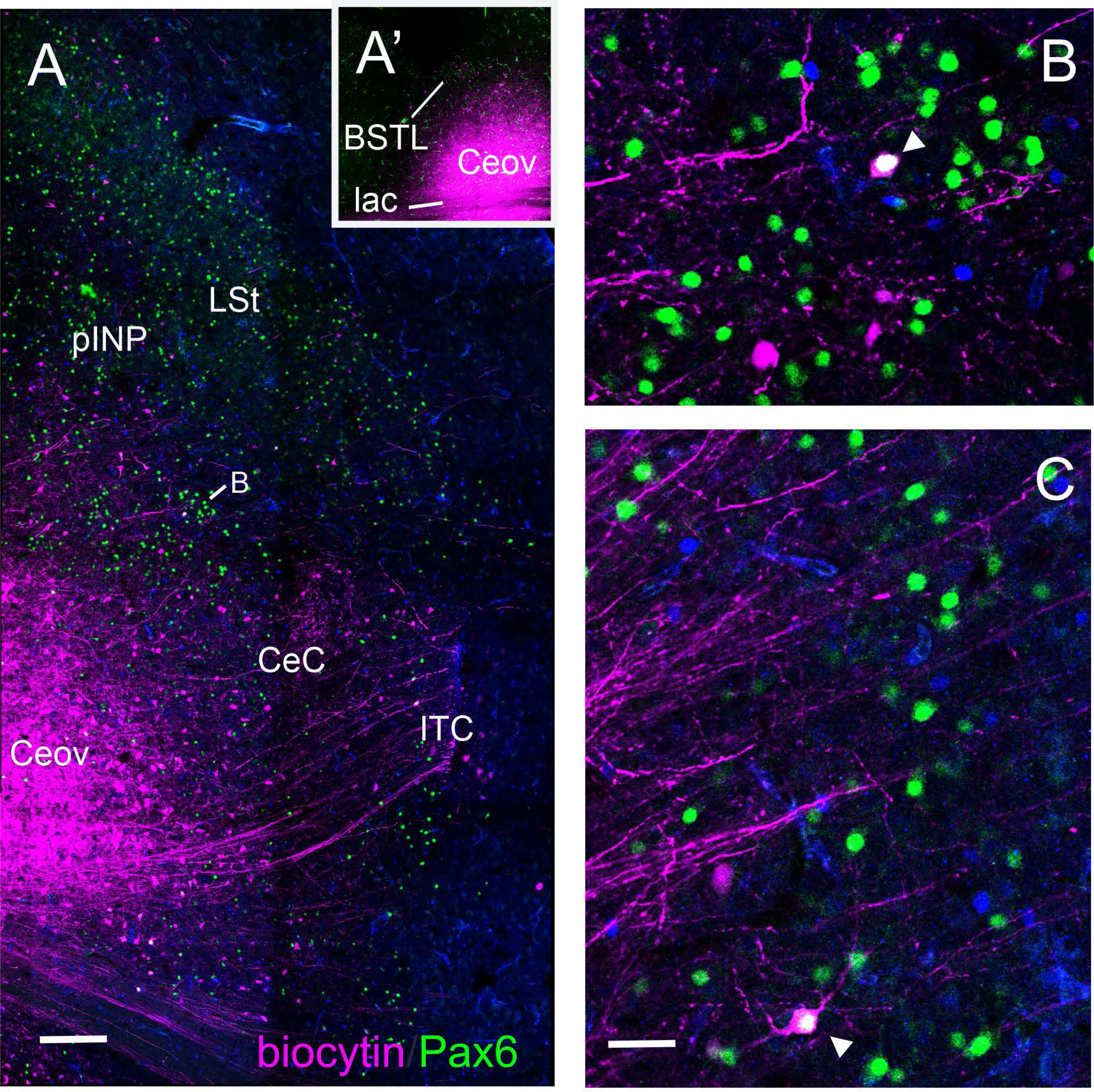
Results of a tract-tracing experiment, following deposit of biocytin crystal in a E18 chicken Ceov at the level of the anterior commissure. A’ shows the location of the biocytin crystal, and A shows retrogradely labeled cells in different subdivisions of the central extended amygdala, including the intercalated amygdala cells (ITC), and the capsular central amygdala (CeC). Double labeling of biocytin with Pax6 showed coexpression in some cells of the medial extension of CeC (detail in panel B) and ITC (detail in panel C) that project to Ceov. See text for more details. For abbreviations see list. Scale bars: A = 500 μm; C = 62,5 μm (applies to B, C).

## Discussion

### Subtypes of CRF cells in the chicken central extended amygdala based on coexpression with different developmental regulatory transcription factors

Our results show the existence of two major groups of CRF cells in the BSTL and central amygdala (mostly located in the medial part of CeC [CeCm], spreading over Ceov) of chicken, resembling those found in mammals. The presence of CRF cells in BSTL agrees with previous findings based on immunohistochemistry in different avian species (chicken and quail: Richard et al., 2004; pigeon: it was identified as nucleus accumbens by Peczély and Antoni, 1984) and in situ hybridization (chicken: Metwalli et al., 2023), while those in the central amygdala were recently found in chicken using in situ hybridization (Metwalli et al., 2023).

Like in chicken, in mammals there are two major groups of CRF cells located in the BSTL and the lateral subdivision of the central amygdala, although these include several subpopulations, based on coexpression with other neuropeptides. A subpopulation of CRF cells in both BSTL and central amygdala coexpresses at least tachykinin-related and neurotensin neuropeptides (Shimada et al., 1989; McCullough et al., 2018; Kovner et al., 2019). The function of these different cell subpopulations is unknown. Since the embryonic origin determines much of the adult phenotype of neurons (Medina et al., 2017, 2023), knowing more on the coexpression of developmental transcription factors in CRF cells of the central extended amygdala in mammals and sauropsids could help to understand other aspects on the molecular profile and connections of these cells, and this information is critical for studying their function. However, coexpression of CRF with developmental transcription factors has not been done in mammals. Based on comparison of expression patterns of CRF with the different developmental transcription factors, it was suggested that CRF cells of the central amygdala might coexpress Islet1 (Bupesh et al., 2011). Our results in chicken point to the existence of at least two subpopulations, one coexpressing Islet1 (which seems relatively more abundant) and another one coexpressing Pax6. This opens the venue to investigate if CRF cells of the mammalian central extended amygdala also include subpopulations with different embryonic origins.

By way of double and triple labeling of CRF with different transcription factors, we found that most CRF cells of chicken BSTL coexpress Islet1 and a few express Pax6. CRF cells coexpressing Islet1 or Pax6 were also found in the medial CeC. Regarding those in BSTL, the major cell population expressing Islet1 may include cells with ventral striatal embryonic origin as well as a few with preoptic origin (Vicario et al., 2015), while the few CRF cells expressing Pax6 likely originate in the dorsal striatal embryonic domain. We exclude an extratelencephalic origin of CRF cells of BSTL because our results show that all of them express Foxg1, a transcription factor involved in specification and subsequent development of the telencephalon (Dou et al., 1999; Martynoga et al., 2005; Manuel et al., 2010, 2011; Metwalli et al., 2022; Medina et al., 2023). Similarly, all the CRF cells of medial CeC (including its medial extension over Ceov) coexpress Foxg1 and have a telencephalic origin, and those coexpressing Pax6 or Islet1 likely originate in the dorsal or ventral striatal embryonic domains, respectively. However, the majority did not coexpress Pax6, Islet1 or Nkx2.1. This may be due to downregulation of the expression of these developmental regulatory transcription factors when the cells begin differentiation and maturation, at late embryonic and peri-hatching stages. Partial downregulation of Nkx2.1 has been observed in the subpallium of late chicken embryos (Pross et al., 2022), and we cannot discard that this also occurs with Pax6 and Islet1, although this seems to be minor compared to that affecting Nkx2.1.

Coexpression of CRF and different developmental regulatory transcription factors could also help to identify homologous cell subpopulations in different species, and may provide insights into their evolution. However, while CRF cells appear to be present in the BSTL of different amniotes, including reptiles (López-Avalos et al., 1993), suggesting that they were present in stem amniotes, data on coexpression with developmental transcription factors is missing in mammals and reptiles. Moreover, CRF cells have not been found in the central amygdala of reptiles (Mancera et al., 1991; López-Avalos et al., 1993; Kavelik, 2021), raising doubts on the homology of those present in mammals and birds. However, the studies on CRF expression in reptiles were done using immunohistochemistry, but it would be convenient to re-evaluate the presence or absence of CRF cells in the central amygdala of reptiles using in situ hybridization, since this technique is more sensitive to detect neuropeptidergic perikarya (as discussed by Metwalli et al., 2023).

### Connectivity between subdivisions of the central extended amygdala in chicken

In our tract-tracing experiments, we found that several subdivisions of the chicken central amygdala project to BSTL, including pINP, CeC and Ceov. This agrees with previous findings in chicken by Hanics et al. (2017), although these authors did not identify which specific subdivisions of the extended amygdala project to BSTL. We also observed labeled fibers in the areas with retrogradely labeled cells, which may include processes of the labeled cells, anterograde labeling of axons from BSTL, as well as fibers of passage (to and from the arcopallium, as shown previously; Atoji et al., 2006; Hanics et al., 2017). Here we focused our attention on the retrogradely labeled cells. The projections of the chicken central amygdala resemble the projections from several subdivisions of the central amygdala to the BSTL in mammals (Dong et al., 2001a). In both mammals and birds, this includes the subdivision that contains the majority of the CRF cells (mammals: the lateral subnucleus; Davis et al., 2010; chicken: present results). Our data on double-labeling of biocytin and different transcription factors showed that some of the retrogradely labeled cells of pINP and CeC coexpress Pax6, while some of those in Ceov coexpress Islet1. However, the majority of the retrogradely labeled cells of the tract-tracing experiments did not show coexpression with any of the transcription factors analyzed here. Since the experiments were done in ex-vivo conditions, perhaps the preservation of the tissue was not perfect leading to very low, undetectable levels of expression of the transcription factors in part of the cells. For this same reason, in situ hybridization for CRF did not work in this tissue, and we could not do double-labeling of biocytin with CRF.

Regarding the Pax6 cells, a previous study showed that a part of them in pINP and CeC coexpresses ENK (Pross et al., 2022), and the present results show that another part coexpresses CRF. This suggests that perhaps both CRF and ENK cells of the chicken central amygdala project to BSTL. This is quite similar to the findings in mammals (Rao et al., 1987; Partridge et al., 2016; Asok et al., 2018). In mammals, CRF and ENK cells of the central amygdala play different roles: while CRF cells are involved in long-term components of fear learning and recall (related to anxiety; Pitts et al., 2009; Davis et al., 2010; Gafford and Ressler, 2015; Asok et al., 2018; Pomrenze et al., 2019), ENK/PKCδ cells promote anxiolysis and analgesia (Paretkar and Dimitrov, 2019; Douceau et al., 2022). Like in mammals, ENK cells and CRF cells of the avian central amygdala have subpallial origin (Vicario et al., 2014, 2015, 2017; Pross et al., 2022; present results) and are likely GABAergic, as typical for cells in the subpallium (Abellán and Medina, 2009; Kuenzel et al., 2011; Medina et al., 2011, 2017, 2023). Thus, projections from these cells to BSTL would lead to inhibition of BSTL cells (Figure 10, scheme). Like in mammals (Gray and Magnuson, 1987, 1992; Moga et al., 1989; Moga and Saper, 1994; Dong et al., 2001b; Choi et al., 2007; Ulrich-Lai and Herman, 2009; Calhoon and Tye, 2015; Zhan et al., 2021), avian BSTL has outputs to the paraventricular hypothalamic nucleus (thus regulating HPA axis), as well as to autonomic and behavior control centers in the hypothalamus and brainstem, such as the parabrachial nucleus, the solitary nucleus and the periaqueductal gray (Atoji et al., 2006; Bálint et al., 2011). Since the avian BSTL contains different types of GABAergic neurons (Vicario et al., 2014, 2015, 2017; Pross et al., 2022; present results), the inhibitory projections from ENK and CRF cells of central amygdala to BSTL might lead to disinhibition of HPA axis and other targets of BSTL, just like in mammals (Ulrich-Lai and Herman, 2009; Zhang et al., 2021). In mammals, CRF cells of BSTL are involved in part of the projections to paraventricular hypothalamus (Moga and Saper, 1994), while the CRF, SST, tachykinin (substance P), and neurotensin cells of BSTL project to autonomic and behavior-control centers in the brainstem (Moga et al., 1989; Gray and Magnuson, 1992). This may be similar in birds. As BSTL of mammals (Csáki et al., 2000; Kudo et al., 2012), the avian BSTL also contains a minor subpopulation of glutamatergic neurons (Abellán et al., 2009; Metwalli et al., 2022). Like in mammals (Ulrich-Lai and Herman, 2009; Kim et al., 2013; Calhoon and Tye, 2015), activation of the latter cells in avian BSTL by inputs from the pallial amygdala (Davies et al., 1997; Dubbeldam et al., 1997; Atoji et al., 2006) could lead to activation of HPA or other targets of BSTL (such as the ventral tegmental area, involved in both reward and aversion; Riters et al., 2012; Jennings et al., 2013; Holy and Miczek, 2016), but their inhibition by GABAergic inputs from the central amygdala or from other cells of BSTL could have the opposite effects. More studies are needed to know the exact connectivity of the ENK and CRF cells of the avian BSTL and central amygdala, and their roles in learning, anxiety and analgesia.

**Figure 10.**
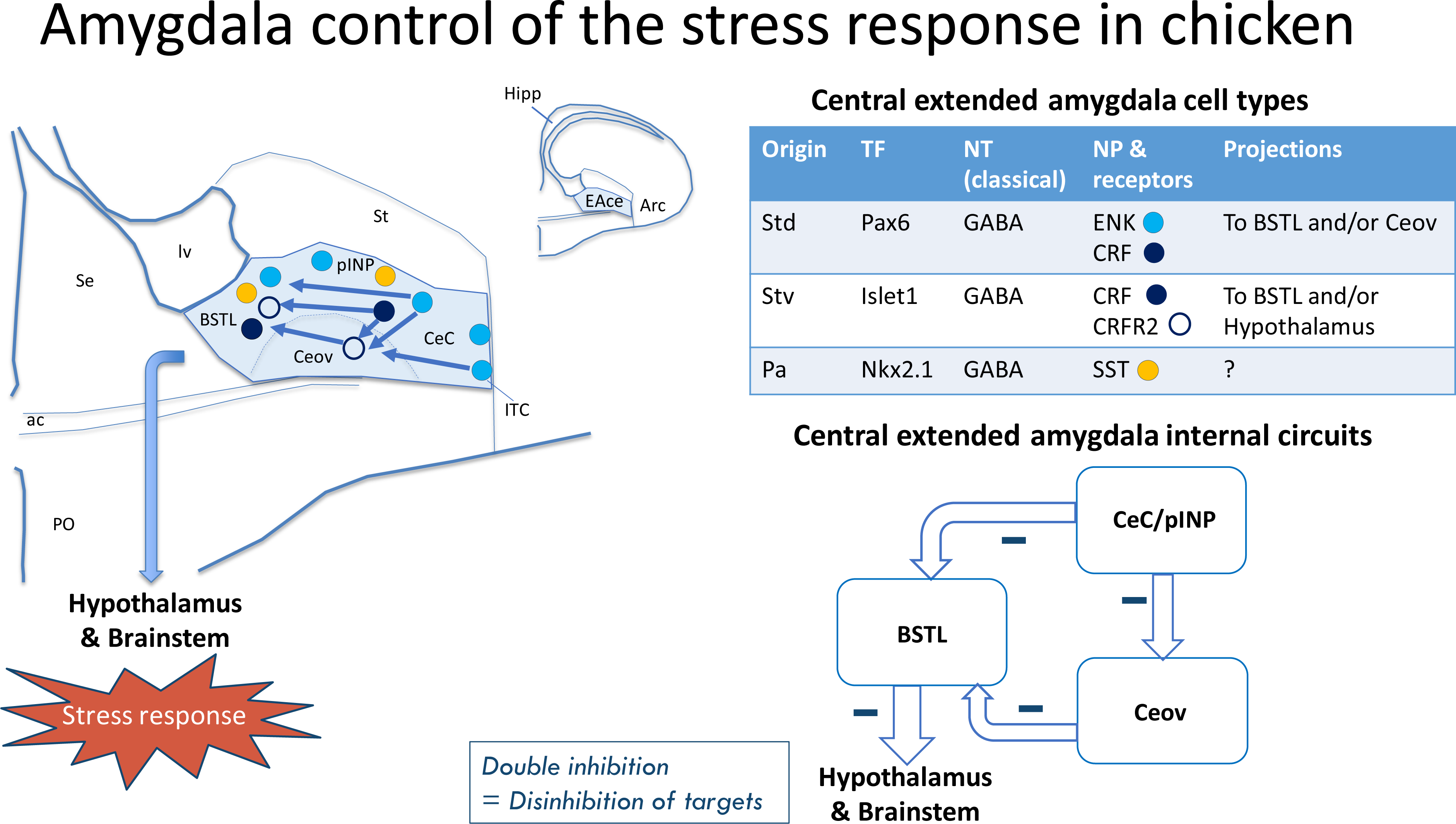
Scheme with a representation of the chicken central extended amygdala, with its subdivisions and some of the major cell types and internal connections found in this and previous studies. CeC and pINP subdivisions (both including Pax6 cells) as well as Ceov subdivision (including Islet1 cells) project directly to BSTL. Since CeC and pINP also appear to project to Ceov, the connections with BSTL can also be indirect by way of Ceov. All of these projections are likely GABAergic, and may involve CRF, ENK and other peptidergic cells. Like in mammals, BSTL contains mainly GABAergic cells and has descending projections to neuroendocrine, autonomic and behavior control centers of the hypothalamus and brainstem, involved in the stress response. Inhibition of BSTL cells by inputs of the central amygdala would lead to disinhibition of the targets, thus producing the stress response. See text for more details. For abbreviations see list.

Regarding the Islet1 cells of Ceov projecting to BSTL, these may establish an indirect link between CeC, pINP and ITC with BSTL (as part of CeC/pINP/ITC-Ceov-BSTL pathways), since our experiments with biocytin in Ceov resulted in retrogradely labeled cells in these three areas. However, this should be confirmed by injecting these areas with an anterograde tracer. CeC and pINP contain ENK and CRF cells, and both might give rise to projections to Ceov, in addition to the projections to BSTL mentioned above. Both BSTL and the area encompassing Ceov express opioid receptors in pigeon (mainly delta and kappa; Reiner et al., 1989), and both also show expression of CRF receptors 1 and 2 (Vicario et al., 2014; Metwalli et al., 2023). Our results on double labeling also show that most Islet1 cells of Ceov express at least CRF receptor 2. These receptors could be mediating the transmission of a projection from CRF cells (at least those expressing Pax6) of CeC and pINP to Islet1 cells of Ceov, which in turn project to BSTL. As noted above, CRF cells of medial CeC and pINP might also project directly to BSTL and, among other targets, might contact Islet1 cells of BSTL that express CRF receptor 2 (present results). Thus, CRF cells of chicken CeC/pINP might influence BSTL both directly as well as indirectly by way of Islet1 cells of Ceov (Figure 10, scheme). This might be similar for the ENK of the chicken CeC/pINP. A previous study showed that Islet1 cells of Ceov also project to the lateral hypothalamus (Vicario et al., 2014). This means that Ceov, including its Islet1 cells, may also play a role in regulation of autonomic related subdivisions of the hypothalamus, as described for the central amygdala of mammals (Swanson and Petrovich, 1998). In chicken, inhibitory projections from CeC, pINP and other central extended amygdala subdivisions to Ceov might lead to disinhibition of the lateral hypothalamus and other putative Ceov targets.

Overall, the results show the existence of complex networks for regulating the physiological and behavioral aspects of stress in chicken, involving different subdivisions and cell types, and open new venues for studying the specific function of different cells and circuits. This information would help to better understand brain regulation of stress in chicken and could be used to improve poultry welfare.

## Author contributions

ED and LM first designed the project, and AP contributed to refine it. AP processed most of the material as part of her Ph.D. research project, and AA and AHM contributed in the processing. AP photographed and prepared many study figures, and analyzed the material with help of LM and ED. AP and LM prepared the figures for the article. AP and LM produced the first draft of the manuscript, ED contributed to improve it, and all authors revised it and approved it.

## Acknowledgments

We deeply thank all Agencies that funded our research. We also thank the technicians and other staff of the Department of Experimental Medicine.

## List of abbreviations

ac: Anterior commissure
af: Amygdalofugal tract
Arc: Arcopallium
BMC: Basal magnocellular corticopetal complex
BST: Bed nucleus of the stria terminalis
BSTL: Lateral BST
BSTLd: Dorsal division of BSTL
BSTLv: Ventral division of BSTL
BSTM: Medial BST
CeC: Capsular central amygdala
CeCl: Lateral part of CeC
CeCm: Medial part of CeC
Ceov: Oval central amygdala nucleus
CSt: Caudal striatum
EA: Extended amygdala
EAce: Central EA
Hipp: Hippocampal formation
ITC: Intercalated amygdalar cells
L: Auditory field L
lac: Lateral limb of ac
lfb: Lateral forebrain bundle
LSt: Lateral Striatum
lv: Lateral ventricle
Me: Medial amygdala
MSt: Medial striatum
Pa: Pallidal embryonic origin
pINP: Peri/post-INP island field
PO: Preoptic region
Pov: Perioval zone
Se: Septum
SpAr: Subpallial amygdala, rostral part
StC: Striatal capsule
Std: Dorsal striatal embryonic origin
Stv: Ventral striatal embryonic origin
vaf: ventral amygdalofugal tract

